# Met-Vision stratifies single cell metabolism revealing coexisting energetic states in tissue macrophages redistributed by inflammation

**DOI:** 10.1101/2025.10.10.681557

**Authors:** Adriana Lecourieux, Margot Bardou, Zacarias Garcia, Philippe Bousso

**Affiliations:** Dynamics of Immune Responses Unit, Institut Pasteur, Université de Paris Cité, INSERM U1223, F-75015 Paris, France

## Abstract

The function of tissue-associated macrophages is tightly linked to their energy metabolism. Yet, the diversity of macrophage metabolic profiles coexisting in tissues at homeostasis or during immune challenges is incompletely understood. Here, we introduce Met-Vision, an imaging-based pipeline for single-cell functional profiling and classification of energy metabolism. Across multiple tissue contexts, we identified that macrophages do not adopt a uniform metabolic profile but typically co-exist in four discrete metabolic states with distinct dependence on OXPHOS and metabolic plasticity. Inflammation reconfigured the distribution of macrophage metabolic profiles that remained heterogeneous. Notably, inflammation-derived nitric oxide finely tuned the distribution of macrophage energetic states. These findings challenge the view of homogenous metabolic activation and reveal a layer of metabolic diversity in tissue at steady state and during inflammation. The ability to stratify macrophage energy metabolic profiles with Met-Vision should help guide the development of metabolism-targeted therapies for inflammatory diseases, cancer, and metabolic disorders.

**Significance statement:** This study introduces *Met-Vision* and reveal that tissue macrophages occupy four distinct energetic states, challenging the notion of uniform metabolic activation and opening avenues for guiding metabolism-targeted therapies.

## Introduction

Macrophages are versatile innate immune cells that contribute to tissue homeostasis, host defense and inflammation (*1, 2*). Their functions are intimately shaped by cellular metabolism, which fuels basic bioenergetic needs and influences key processes such as cytokine production, phagocytosis, antigen presentation, and cell survival (*3–5*). Over the past decade, a growing body of work has supported the idea that metabolic reprogramming is a central driver of macrophage identity and function (*6, 7*) in infection and cancer (*8, 9*).

A widely accepted framework suggests that resting or alternatively activated macrophages rely primarily on oxidative phosphorylation (OXPHOS) for energy production. By contrast, macrophages switch to aerobic glycolysis in response to classical activation with pro-inflammatory stimuli such as LPS and IFN-γ (*10–13*). This model, largely derived from *in vitro* studies, has provided valuable insights into macrophage polarization and its link to immunometabolism.

Emerging evidence from single cell transcriptomic, proteomic, or metabolomic studies suggests that macrophage metabolism is more heterogeneous than previously thought (*14–19*). Most of these studies however infer energy metabolism from indirect readouts (gene or protein expression programs), which may incompletely reflect functional energy metabolism at the single cell level. Alternatively, functional measurements can be classically performed using extracellular metabolic flux assays, but this important approach lacks single cell resolution. Additionally, the SCENITH technology (*20*) that measures protein synthesis as a proxy for ATP is a powerful to approach to analyze rare immune cell populations and can be combined with other read-outs to profile macrophage immunometabolism up to the subpopulation level (*21*). Therefore, despite the existence of many complementary methodologies, the diversity of macrophage metabolic profiles coexisting in native tissue environments and its modulation by inflammation remain incompletely understood.

To functionally profile the energy metabolism of individual macrophages *ex vivo*, we have recently introduced SPICE-Met (*22*), an approach relying on a new transgenic mouse (termed SPICY mouse) expressing a fluorescent reporter for cellular ATP:ADP ratio (*23*). Immune cells from SPICY mice can be challenged with metabolic drugs to analyze their dependence on specific metabolic pathways using flow cytometry or imaging (*22*). While SPICE-Met provides valuable functional information on energy metabolism, it is currently constrained by a relatively low throughput and reliance on manual analysis. To overcome these limitations, we introduce here Met-Vision, a computer vision-based pipeline designed to extract, interpret and classify functional metabolic information from live-cell imaging data with single cell resolution.

Using this approach, we uncover the presence of four discrete metabolic states within macrophage populations across tissues. Moreover, we show that inflammatory cues shift the distribution of these states rather than inducing a uniform metabolic reprogramming. This study provides a novel framework to investigate tissue macrophage bioenergetics in health and disease.

## Results

### Imaging macrophage energy metabolism at the single cell level

To functionally profile energy metabolism in individual macrophages, we developed Met-Vision, a fully automated, computer vision–based pipeline for single-cell metabolic profiling. Met-Vision builds on our previous SPICE-Met approach (*22*), introducing scalability and machine-learning based classification to high-throughput image analysis. The method uses SPICY mice (*22*), a transgenic line expressing the fluorescent ATP:ADP sensor PercevalHR (*23*) in immune cells. Macrophages isolated from SPICY mice were imaged before and after treatment with oligomycin, an inhibitor of mitochondrial respiration (**Figure 1A**). Met-Vision tracks dynamic changes in ATP:ADP ratio over time in hundreds of individual cells. Following segmentation and normalization, the pipeline generates time-lapse arrays of single-cell responses (**Figure 1A-B**), which can also be visualized as kinetics traces (**Figure 1C**) or heatmaps (**Figure 1D**) of normalized ATP:ADP ratios, capturing energetic status before and after OXPHOS inhibition.

**Figure 1:**
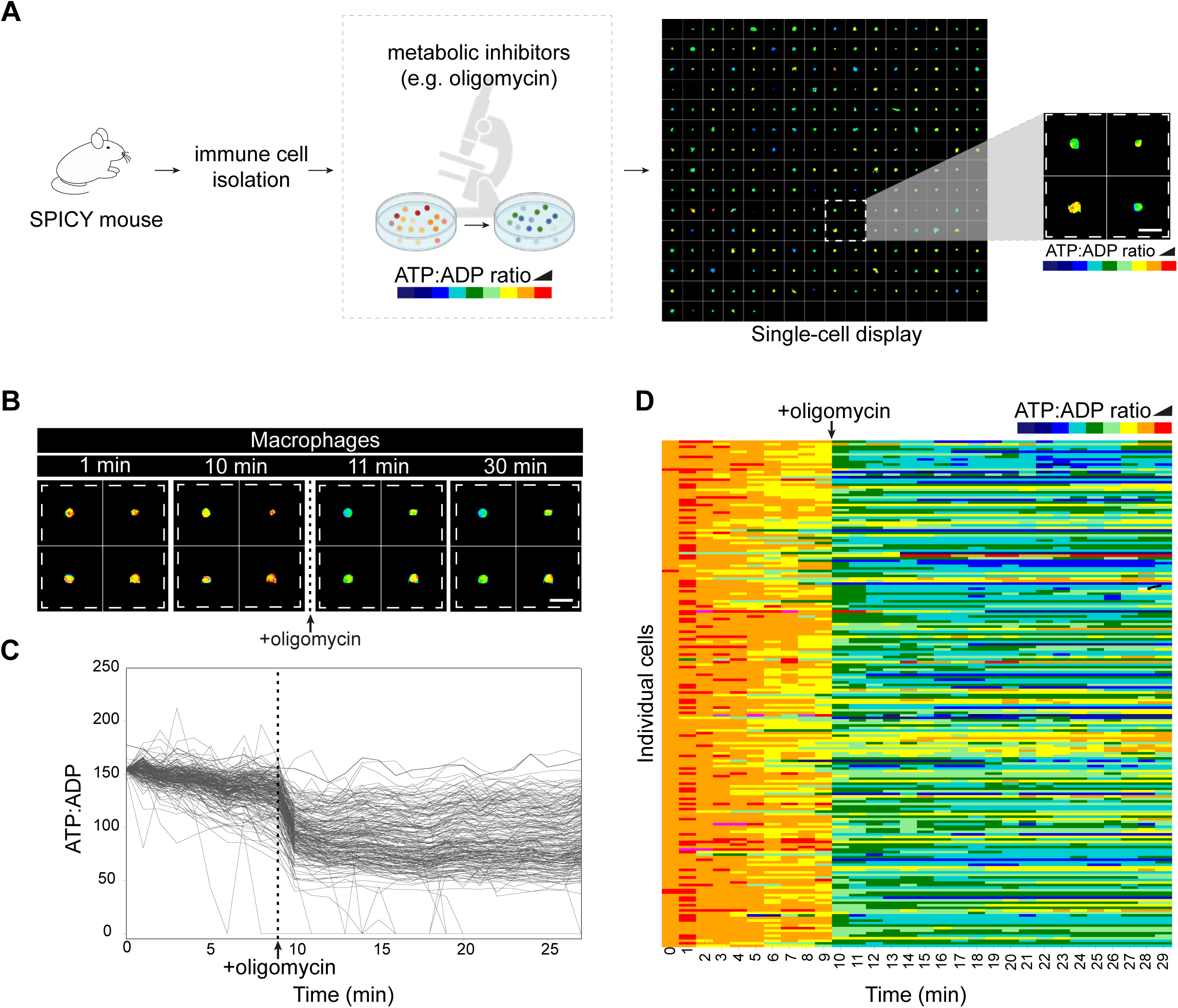
Met-Vision, an imaging-based automated pipeline to profile energy metabolism at the single-cell level. **A)** Principle for Met-Vision analyses. Cells isolated from SPICY mice expressing the PercevalHR fluorescent reporter for ATP:ADP are imaged before and after addition of oligomycin. After cell segmentation and normalization steps, the Met-Vision pipeline extracts images and values for single cells. Met-Vision generates a single-cell time-lapse array of response of ATP:ADP dynamics. It is expected that cells with strong dependence on OXPHOS for energy production exhibit a rapid drop in ATP:ADP ratio in response to oligomycin while cells primarily dependent on glycolysis maintained a stable energetic status. **B**) Example of single macrophage responses to oligomycin. **C-D**) The dynamics of ATP:ADP ratio for individual cells is represented as curves (**C**) or heatmaps (**D**). Scale bars represent 30 µm.

### A classifier for macrophage energy metabolic profiles

To further describe the diversity of macrophage metabolism within a given sample, we sought to estimate the number of distinct energy metabolic profiles in macrophages. We generated a large dataset of single cell responses (**Figure 2A**), compiling imaging of bone marrow derived macrophages (BMDMs) and peritoneal macrophages treated or not with an inflammatory stimulus (LPS and IFN-γ). Using Kmeans clustering, we identified 4 distinct types of macrophage response to oligomycin treatment (**Figure 2B**). Cells in the first cluster corresponded to cells with maximal decrease in ATP:ADP upon oligomycin treatment, reflecting maximal dependence of OXPHOS (Ox-high). The second cluster contained cells that exhibited a partial but sustained decrease in energetic status suggesting a concomitant use of OXPHOS and glycolysis (Ox-med). Macrophages in the third cluster displayed a small decrease in ATP:ADP ratio that was rapidly reversed suggesting a plastic metabolic phenotype with the capacity to rapidly compensate for the drop in energetic resources through glycolysis (Ox-low). Finally, the fourth cluster corresponded to cells that appeared largely independent on OXPHOS and were strictly dependent on glycolysis (Ox-null).

**Figure 2:**
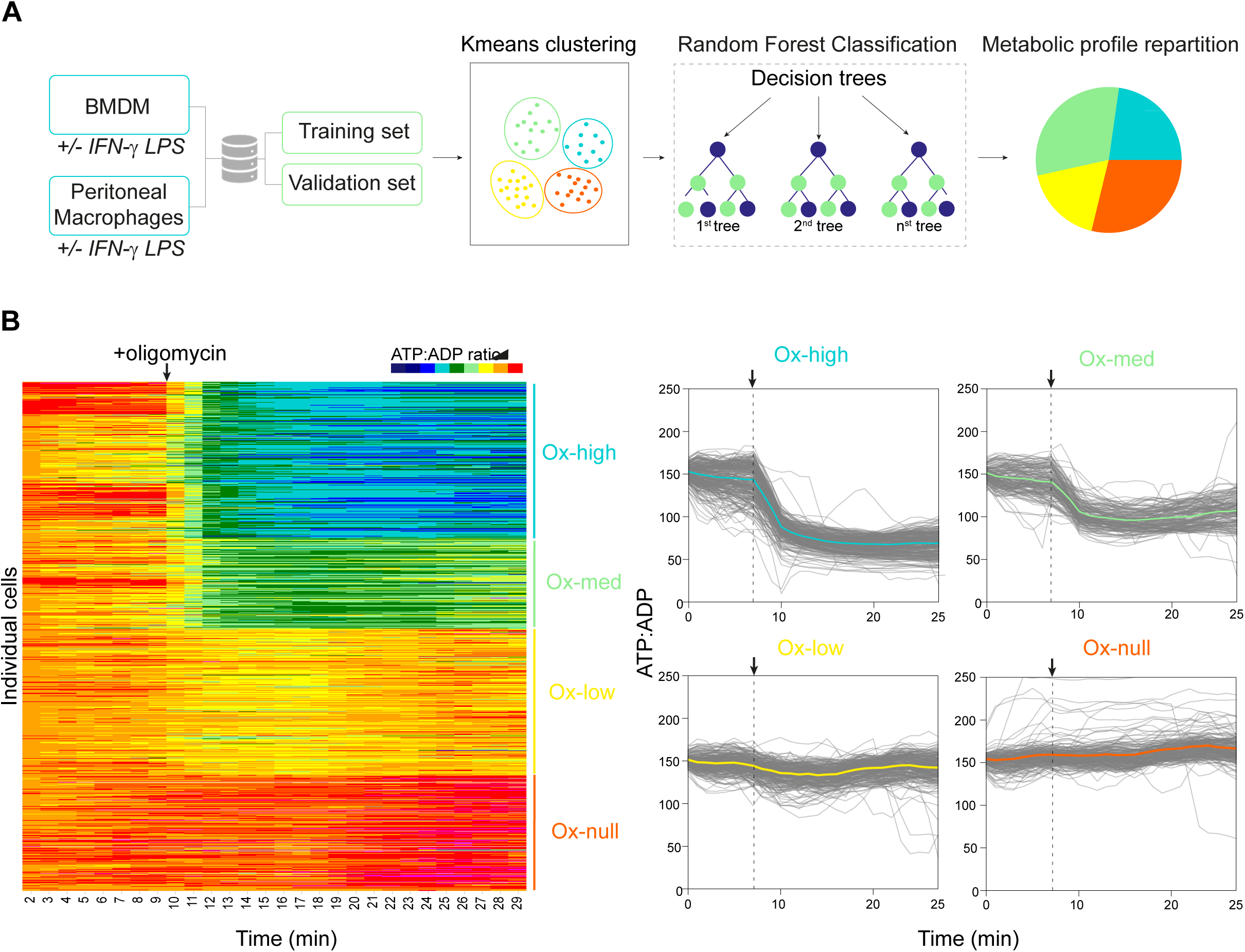
Met-Vision identifies and classifies distinct energy metabolism profiles in macrophages. **A)** Scheme illustrating the main steps to establish a classifier for macrophage energy metabolic profiles. The dataset (n=879 macrophages) containing kinetic traces of ATP:ADP ratios for individual macrophages treated with oligomycin was built from resting or activated (LPS+IFN-γ) BMDM and peritoneal macrophages. For each condition, data are from at least 2 independent experiments with 2-3 imaging fields analyzed in each experiment. Four metabolic profiles were defined using the k-means clustering algorithm. The clustered dataset was divided into a training set (80%) and a validation set (20%) to build a Time Series Forest classifier. Met-Vision outputs the distribution of metabolic profile within the imaged population in the form of a color-coded pie chart. **B**) Heatmaps (left) and kinetic curves (right) showing the response of individual macrophages to oligomycin after clustering. Black arrows correspond to oligomycin addition.

In order to identify the distribution of these four energy metabolic profiles within a given cell population, we trained a classifier model (using time series random forest classification) with a training set comprising n=879 macrophages. When applied to a validation dataset, our model exhibited high accuracy and specificity (**Figure 2A**, **Figure S1**). In sum, the Met-Vision pipeline readily extracts and classifies the diversity of macrophage metabolic profiles in a given cell population.

### Diversity of energy metabolism in macrophages at steady state and upon inflammation

As an initial validation, we applied Met-Vision to BMDMs that were either left untreated or activated with LPS and IFN-γ. In line with previous knowledge, untreated macrophages predominantly exhibited Ox-high and Ox-med metabolic profiles, whereas classically activated macrophages showed a marked shift toward Ox-low and Ox-null states (**Figure 3A, movie 1**). We then extended our analysis to *ex vivo* isolated tissue-associated macrophages, beginning with peritoneal macrophages. At steady state, these cells displayed all four metabolic profiles, with a predominance of Ox-high and Ox-med states (**Figure 3B-C**, **movie 2**). To assess how inflammation reshapes this metabolic landscape, mice were treated *in vivo* with LPS and IFN-γ. Met-Vision analyses revealed a redistribution of metabolic profiles in peritoneal macrophages, now dominated by Ox-low and Ox-null states (**Figure 3B-C**, **movie 2**). One hallmark of inflammatory macrophage activation is the induction of iNOS and subsequent production of nitric oxide. Because nitric oxide has been shown to inhibit mitochondrial respiration (*24, 25*), we asked whether nitric oxide influence the distribution of metabolic profiles. Profiling iNOS deficient (*Nos*2*^-/-^*) SPICY mice, we observed a decrease in the contribution of the Ox-null profile at the expense of Ox-low/Ox-med states in BMDM and *ex vivo* isolated peritoneal macrophages upon inflammation (**Figure 3D-E**). These observations suggest that nitric oxide production finely tunes the distribution of energy metabolic profiles in tissue macrophage during inflammation.

**Figure 3:**
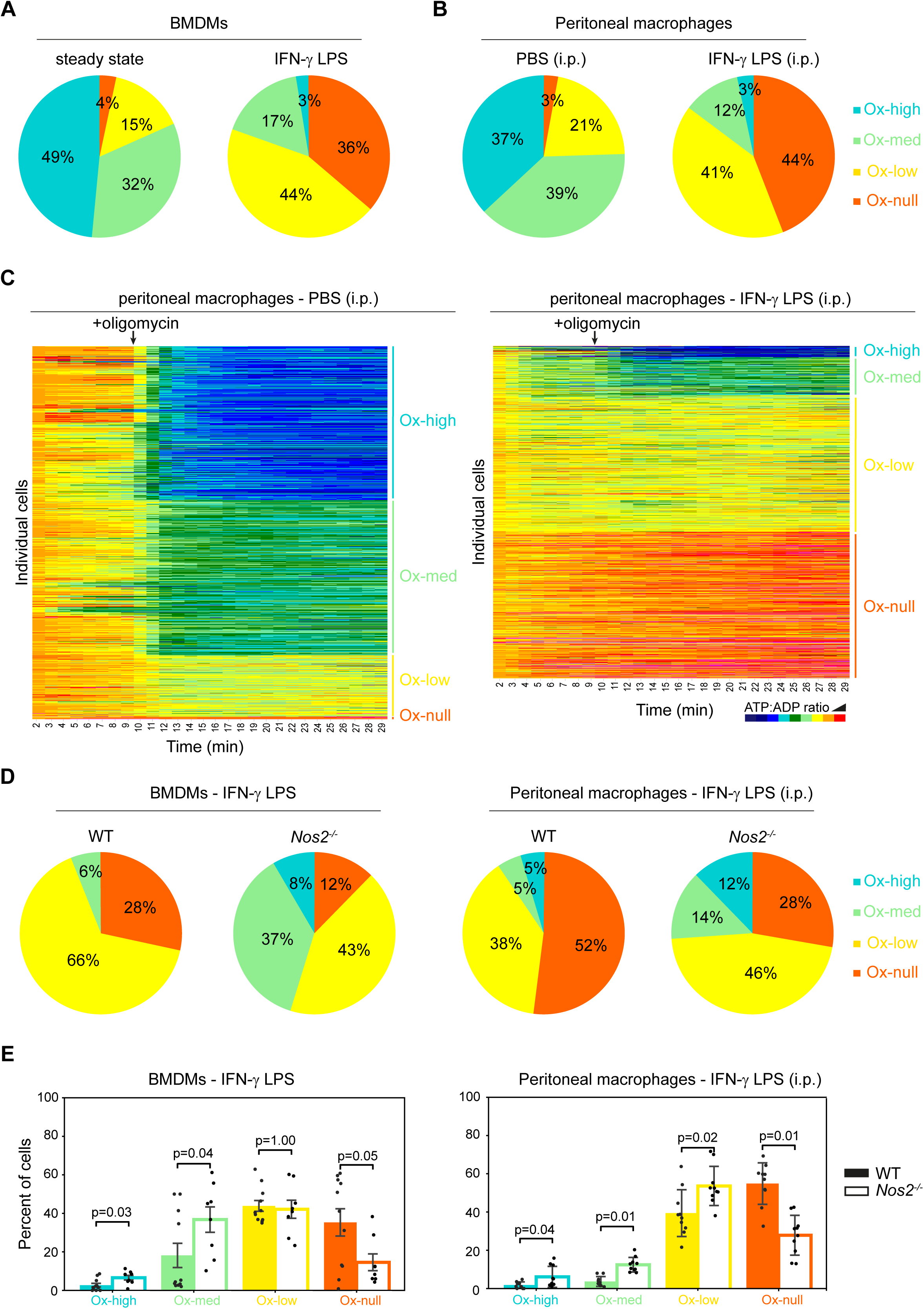
Distinct macrophage metabolic profiles co-exist at steady state and are reconfigured upon inflammation. **A-B**) Distribution of the four metabolic profiles (Ox-high, Ox-Med, Ox-low and Ox-null) in (**A**) BMDM treated or not with LPS+IFN-γ or in (**B**) peritoneal macrophages isolated two days after intraperitoneal (i.p.) injection of LPS and IFN-γ or PBS. Results are compiled from 4-6 independent experiments for each condition with typically 3 distinct imaging positions per experiment. **C**) Heatmaps showing the response of individual peritoneal macrophages to oligomycin classified by profiles at steady state and upon intraperitoneal injection of LPS and IFN-γ. **D-E**) Nitric oxide production fine tunes the distribution of macrophage metabolic profiles in inflammatory conditions. **D**) Distribution of energy metabolic profiles within WT or *Nos2*^-/-^ BMDMs activated with LPS and IFN-γ (left) or within WT or *Nos2*^-/-^ peritoneal macrophages isolated two days after intraperitoneal injection of LPS and IFN-γ (right). Representative of 2 independent experiments. **E**) Quantification of each metabolic profile within (left) WT or *Nos2^-/-^* BMDMs activated with LPS+IFN-γ or (right) peritoneal macrophages isolated from WT or *Nos2^-/-^* SPICY mice injected with LPS+IFN-γ. Each dot represents an imaging position. Compiled from 3 independent experiments with 2-3 imaging positions per experiment. Statistical analyses were performed using a Wilcoxon Mann Whitney test.

Together, these results align with the well-established shift from oxidative phosphorylation to glycolysis during classical activation. However, rather than a uniform metabolic switch, our data uncover a nuanced and dynamic remodeling of energy metabolism, where inflammation and associated nitric oxide production alters the distribution across multiple coexisting metabolic states, revealing heterogeneity that would be masked by population-level analyses.

### Metabolic profiling of macrophages with distinct phenotypes using Met-Vision

Several mechanisms could account for the overall diversity of energy metabolic profiles including differences in ontogeny and/or phenotype. We reasoned that integrating phenotypic marker analysis with functional metabolic profiling could offer deeper biological insights. To this end, we developed the ability to simultaneously image the cellular ATP:ADP ratio alongside two cell surface markers, labeled with fluorescently conjugated monoclonal antibodies, enabling concurrent assessment of metabolism and phenotype at the single-cell level. As a proof of concept, we analyzed peritoneal macrophages, which comprise two well-characterized subsets: resident macrophages, expressing high levels of F4/80 and low levels of MHC class II, and monocyte-derived macrophages, which show the opposite profile. As previously described, these subsets can be readily distinguished by flow cytometry using these two surface markers (**Figure 4A**). To enable similar resolution in our imaging-based approach, cells were stained with fluorescent antibodies against F4/80 and MHC class II prior to image acquisition. Met-Vision allows for the stratification of metabolic profiles based on marker expression using a gating strategy analogous to flow cytometry. As shown in **Figure 4B**, the two macrophage subsets were clearly identified by Met-Vision. Importantly, subsequent metabolic analysis revealed relatively equivalent distributions of the four metabolic profiles for resident and monocyte-derived macrophages at steady state (**Figure 4C**). These results illustrate the possibility to further stratify macrophage metabolism within defined subpopulation.

**Figure 4:**
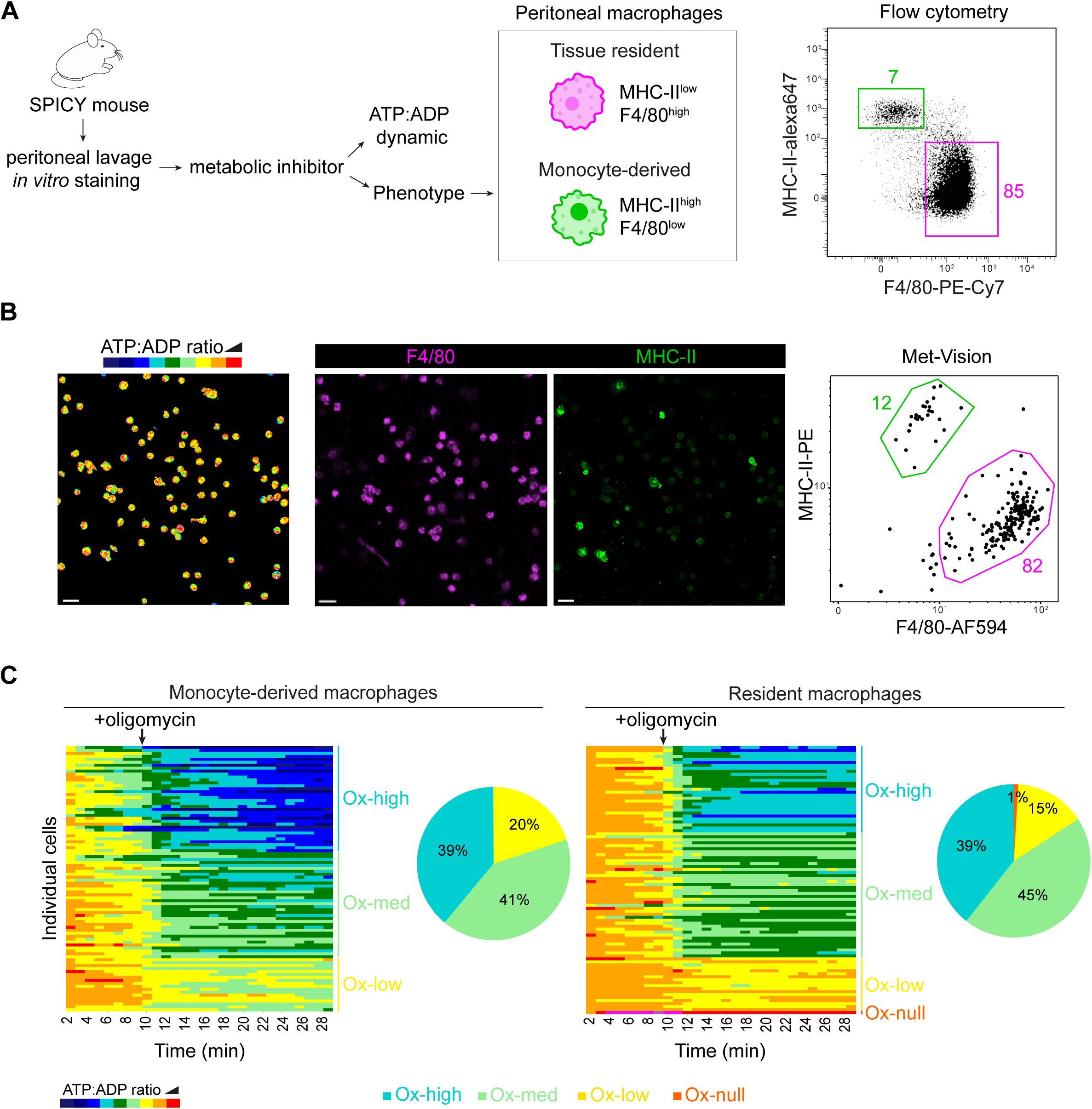
Linking macrophage phenotypes and metabolic profiles using Met-Vision. **A**) Peritoneal macrophages classically comprise two populations, resident macrophages and monocyte-derived macrophages easily identified by flow cytometry. **B**) The Met-Vision pipeline integrates optional antibody staining of two surface markers to link phenotype and metabolic profile. Met-Vision represents cell phenotypes in the form of dot plots with the possibility to subdivide metabolic profiling to specific subsets using a gating strategy analogous to flow cytometry. **C**) Heatmaps and distributions of metabolic profiles at steady state in resident and monocyte-derived macrophages. Compiled from 3 independent experiments. Scales bars represent 30 µm.

### Metabolic diversity of alveolar macrophages at steady state and during inflammation

Having established metabolic diversity in peritoneal macrophages, we next turned to lung-resident alveolar macrophages (**Figure 5A-D**) to assess whether metabolic heterogeneity is a generalizable feature of tissue-associated macrophages. Alveolar macrophages are essential for maintaining pulmonary homeostasis and orchestrating responses to inhaled pathogens and environmental insults. We first applied Met-Vision to alveolar macrophages under steady-state conditions. As shown in **Figure 5C, movie 3**, all four energy metabolic profiles were detected, with a predominance of the Ox-high and Ox-med profiles.

**Figure 5:**
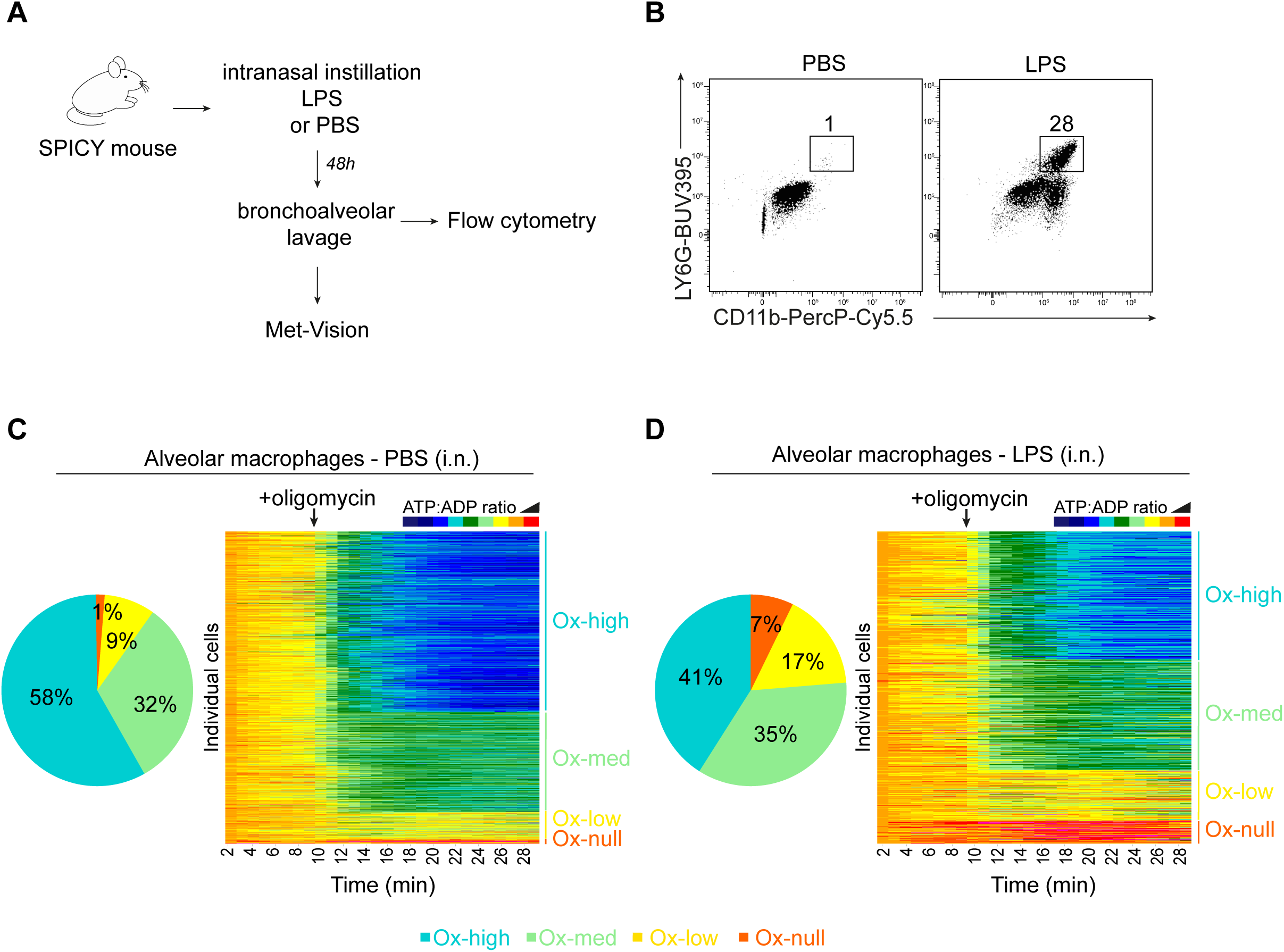
Metabolic profiling of alveolar macrophages at steady-state and during inflammation. **A**) Alveolar macrophages were isolated from SPICY mice that were left untreated or subjected to LPS intranasal (i.n.) instillation two days before. **B**) Flow cytometric profiles showing induction of neutrophilic influx in response to LPS instillation. Representative of 2 independent experiments with 3-4 mice per condition per experiment. **C-D**) Heatmaps and repartition of metabolic profiles within alveolar macrophages at steady-state (C) and upon inflammation (D). Compiled from 3-4 experiments per condition.

To trigger lung inflammation, mice were instilled with LPS. Robust neutrophil recruitment confirmed the initiation of an inflammatory response (**Figure 5B**). Met-Vision profiling revealed only modest changes in the distribution of metabolic profiles, with a moderate increase in the proportion of Ox-low and Ox-null cells, while the Ox-med profile remained largely preserved (**Figure 5D, movie 3**). Mild changes in metabolic profiles compared to untreated mice were also noted when a mixture of LPS and IFN-γ was instilled (**Figure S2**). Unlike peritoneal macrophages, which undergo pronounced metabolic shifts upon inflammation, alveolar macrophages exhibit only a partial and restrained metabolic response to LPS or LPS+IFN-γ instillation, underscoring context-specific differences in metabolic reprogramming by inflammation.

## Discussion

In this study, we functionally profile macrophage energy metabolism using Met-Vision, a new imaging-based pipeline for high-throughput analysis and classification of energy metabolism at single-cell resolution. We revealed discrete metabolic states that coexist within tissues and shift in distribution during inflammation, offering new insights into the dynamic and heterogeneous nature of macrophage metabolism *in vivo*.

Building on original SPICE-Met approach (*22*), Met-Vision integrates real-time imaging of ATP:ADP dynamics with machine learning-based classification to systematically map metabolic states across tissue populations at the single-cell resolution. Additionally, Met-Vision allows simultaneous imaging of cell surface markers, providing the ability to couple metabolic profiling with phenotypic characterization in a manner analogous to flow cytometry but with functional metabolic readouts. Met-Vision can be easily adapted for the study of other immune cell types and should provide a powerful framework for quantitatively dissecting immune metabolism of tissue populations at single-cell resolution.

The classical model of macrophages polarization (*26*) has been associated with a binary framework in which pro-inflammatory (M1) macrophages rely on glycolysis while anti-inflammatory (M2) macrophages rely on OXPHOS. Using Met-Vision, we identified four discrete macrophage metabolic profiles. In addition to metabolic profile corresponding to a full dependence on OXPHOS (Ox-high) or on glycolysis (Ox-null), we identified a profile showing relatively equal contribution of OXPHOS and glycolysis in energy production (Ox-med) as well as a population with a limited but detectable contribution of OXPHOS and strong plasticity (Ox-low). This heterogeneity was evident even at homeostasis, suggesting that macrophage energy metabolism is shaped by more than activation state alone. It is tempting to speculate that a balance of metabolic profiles may be critical for tissue integrity and immune regulation.

We further showed that inflammation does not uniformly reprogram macrophage metabolism but instead shifts the distribution among pre-existing metabolic profiles. This observation again challenges the traditional model of a binary metabolic switch and supports a model of dynamic, subset-specific metabolic remodeling. It has been proposed that macrophage functions are not only dictated by inflammation but also influenced by ontogeny, local cues and duration of tissue residence (*27*). Our results suggest that such complexity is reflected at the level of energy metabolism within a given tissue. Notably, nitric oxide was found to promote the Ox-null energetic profile at the expense of Ox-low/Ox-med states during inflammation suggesting that the level of iNOS induction in tissue fine-tunes the rewiring of energetic states.

Importantly, our data indicate that the extent of energy metabolism reprogramming in macrophage upon inflammation may be tissue-specific and appeared relatively mild in alveolar macrophages, possibly linked to a defect in glycolysis upregulation (*28*).

Recent studies have inferred heterogeneity in macrophage metabolic programs *in vivo* from single-cell transcriptomic or high dimensional flow cytometry datasets. For example, flow cytometry-based profiling of nutrient transporters and metabolic enzymes identified distinct metabolic signatures within tissue-resident macrophages as well as subset-specific responses to infection (*17*). Our results extend these findings by offering a functional perspective on macrophage metabolism directly measuring dependence of cellular energetic resources on metabolic pathways. Future studies using Met-Vision may help refine the relationship between transcriptomic or proteomic signatures and functional energy metabolism.

In sum, by measuring real-time ATP:ADP dynamics, we revealed discrete metabolic states that coexist within tissues and shift in distribution during inflammation, offering new insights into the dynamic and heterogeneous nature of macrophage metabolism *in vivo*. A refined understanding of tissue macrophage bioenergetic diversity will help identify metabolic fingerprints associated with disease states or response to therapy in cancer, inflammatory or metabolic diseases.

## Material and Methods

### Mice

SPICY (Vav-iCre x Perceval^fl/fl^) (*22*) and iNOS KO SPICY (SPICYxNos2^-/-^) mice were bred in the animal facilities of Institut Pasteur. All mice were housed under pathogen-free conditions. All animal studies were approved by the Institut Pasteur Safety Committee in accordance with French and European guidelines (CETEA n°190148 and n°240110).

### *In vitro* BMDM stimulation

Murine bone marrow derived macrophages (BMDMs) were generated as previously described (*25*). Seven days after plating, adherent BMDMs were detached using PBS EDTA 5 mM and seeded into tissue-culture treated dishes at a density of 2×10^5^ cells/cm^2^ with 10% heat-inactivated fetal calf serum (FCS), 100 U/mL penicillin, 100 ng/mL streptomycin, 1 mM sodium pyruvate, 1 mM HEPES and 5 mM 2-mercaptoethanol (complete RPMI) supplemented or not with 1 μg/mL lipopolysaccharide (LPS) (Sigma) and 50 ng/mL IFN-γ (Peprotech) for 24 h.

### *In vivo* treatments

To induce an inflammation in the peritoneal cavity, mice were injected intraperitoneally with a mixture of 25 µg IFN-γ and 25 µg LPS in a total volume of 200 µL PBS. Control mice were injected with PBS only. To induce inflammation in the lung, 10 µg of LPS alone or in combination with 1 µg IFN-γ (in a total volume of 20 µL of PBS) were intranasally instilled in SPICY mice nostrils.

### Isolation of peritoneal and alveolar cells

Forty-eight hours after *in vivo* stimulations, peritoneal and bronchoalveolar lavages were performed. For peritoneal lavages, 8 mL of cold PBS supplemented with 2 mM EDTA and 2% FCS (retrieval buffer) were injected in the mouse abdomen. Following gentle abdominal massage, mice were inverted over 50 mL collection tubes to recover the lavage fluid containing peritoneal macrophages. To collect alveolar macrophages, the lungs were flushed three times with 1 mL of retrieval buffer, which was then aspirated back for cell recovery.

### Cell preparation and staining prior to imaging

Following macrophages isolation, cells were seeded at a density of 2 x 10^5^ cells/cm^2^ in 10 mm glass-bottom poly-L-lysine coated dishes (World Precision Instruments) in 200 µl complete RPMI. Cells were incubated for 4 h at 37°C under 5% CO_2_ to promote macrophages adhesion. After 4 h, non-adherent cells were removed by gentle pipetting of the media. For flow cytometry analysis, adherent peritoneal macrophages were detached using PBS EDTA 5 mM. For live imaging, BMDMs, peritoneal and alveolar macrophages were directly stained in the culture dish for 15 min at 37°C in complete RPMI containing AF594-conjugated anti-mouse F4/80 (clone BM8, BioLegend) and PE-conjugated anti-mouse I-A/I-E (clone M5/114.15.2, BioLegend). After a washing step, cells were placed in RPMI without phenol supplemented with 0.5% FCS and 1% GlutaMAX (Gibco) and kept at 37°C before imaging.

### Flow cytometry

For peritoneal macrophages subset analysis, single cell suspensions were stained for 20 min at 4°C with PE-Cy7-conjugated anti-mouse F4/80 (clone BM8, BioLegend) and Alexa647-conjugated anti-mouse I-A/I-E (clone M5/114.15.2, BioLegend). For alveolar macrophages, the bronchoalveolar lavage cell suspension was stained with BUV395-conjugated anti-mouse Ly6G (clone 1A8, BD Biosciences) and PerCP-Cy5.5 anti-mouse CD11b (clone M1/70, BioLegend). Samples were acquired using a Cytoflex LX (Beckman Coulter) and analyzed using FlowJo software version 10.10.

### ATP:ADP ratio real-time imaging

Visualization of macrophages energy metabolism was performed using an upright Stellaris DIVE two-photon imaging system (Leica), equipped with a Vision 2 laser (Coherent), a galvano scanner and a 25X/1.05 NA objective (Olympus). Excitation was set at 850nm for ADP and AF594-conjugated Abs detection and at 1000nm for ATP and PE-conjugated Abs detection. Detection was performed using the 4Tune module (Leica) from 495 to 565 nm (for ADP and ATP signals), from 570 to 600 nm (for PE-conjugated Ab) and from 640 to 690 nm (for AF594-conjugated Ab). Image stacks with 3 z-planes were collected every minute. Image acquisition was performed for 10 min in the absence of oligomycin and continued for 20 min after the addition of 1 µM oligomycin.

### Visualization of single cell metabolic dynamics with Met-Vision

To extract and visualize metabolic dynamics in single cells from the raw time-lapse images, we first corrected XY drift in the time-lapse images using the Fast4DReg plugin. Drift-corrected images were then processed with a custom macro written in the ImageJ macro language using the FIJI macro editor. The macro aims to extract time-lapse movies of individual cells and compile these movies into a large cell array. The macro is designed to analyze time-lapse images containing 2 to 4 channels. Two of them correspond to ATP and ADP fluorescence, while the additional channels can be used to incorporate cell surface markers into the analysis. Briefly, a ratio image was first generated by performing a pixel-wise calculation of the intensity ratio between the ATP and ADP channels ((ATP channel / ADP channel) × 100). Region of interests (ROIs) were segmented from the red (ADP) channel using Li’s thresholding method, followed by morphological processing and watershed to refine object detection. A size filter was applied to retain particles larger than 50 pixels. A time-lapse cell array is generated, where each ROI is placed into a 100×100-pixel tile and arranged into a square grid. An optional normalization step adjusts the relative intensity of each ROI by scaling to a target mean intensity of 155 on frame 1. The final output is visualized using a 16-color lookup table. Based on user-defined settings, the time-lapse cell array together with individual tiles are saved in TIFF format.

### Met-Vision analysis pipeline for classification of energy metabolic profiles

To analyze and classify energy metabolic profiles, we established a homemade pipeline using Python version 3.11.5. For each individual tile generated by the ImageJ macro, the script calculated the mean intensity values of the ATP:ADP ratio over time. Subsequent steps included normalization, missing values imputation, and smoothing with a rolling mean (window = 3). Cells with ATP:ADP ratios containing zeros or values below 50 in the first 8 time points were considered leaving the field of view or incorrectly identified and were excluded from the analysis. The dataset generated is consolidated into a single CSV file containing the ATP:ADP ratio values for each ROI at every time point, along with the intensity values of other markers at frame 1. The pipeline next assigns each cell from the CSV file to a class based on its metabolic profile dynamics. Metabolic classes were defined using the k-means clustering algorithm, applied to data from 8 movies comprising a total of 879 individual cells, resulting in the identification of four distinct metabolic clusters. Clustering was tested with *k* values ranging from 3 to 6. Based on both the elbow method and the biological interpretability of the resulting clusters, the choice of *k* = 4 was retained. The clustered dataset was partitioned into a training set (80%) and a validation set (20%) for the construction of a Time Series Forest classifier, implemented with the `sktime` library using default parameters. To evaluate the performance of the model we used several metrics:

- Precision (Positive Predictive Value) = TP/(TP+FP)
- Recall (True Positive Rate, Sensitivity or Coverage) = TP/(TP + FN)
- F1-score = 2 × (Precision × Recall)/(Precision + Recall)

A confusion matrix was computed to provide a detailed overview of classification results across all classes. Receiver Operating Characteristic (ROC) curves were also plotted for each class in a one-vs-rest scheme, and the Area Under the Curve (AUC) was calculated to assess the model’s ability to discriminate between classes.

The model is integrated into the analysis pipeline and automatically generates predictions for each ROI in newly processed movies. Results are visualized as heatmaps, time-series plots with trajectories color-coded by metabolic profiles and pie charts representing the distribution of metabolic profiles in the population. When two additional markers are included in the analysis, a graphical user interface (GUI) implemented with Tkinter displays a FACS-like scatter plot of the markers. The user can delineate subpopulations by drawing polygons directly on the scatter plot. Classification is then performed both on the entire population and independently on each user-defined subpopulation. The pipeline can be tailored through various parameter settings, all of which are detailed in the built-in help section. It is intended to operate via the command line and features an intuitive interface, making it accessible to non-expert users. All the scripts are available in Github (https://github.com/AdrianaBioinfo/perceval/tree/main).

## Acknowledgments

We are grateful to the members of the Bousso laboratory for their valuable feedback on the manuscript. We acknowledge the mouse facility at Institut Pasteur for their support in conducting this study. The work was supported by Institut Pasteur, INSERM and an ANR Grant (Metaniche).

## Author contributions

MB and ZG conducted the experiments; AL designed the bioinformatic pipeline; MB, AL and PB analyzed the experiments and wrote the manuscript.

## Supplementary materials

**Figure S1:**
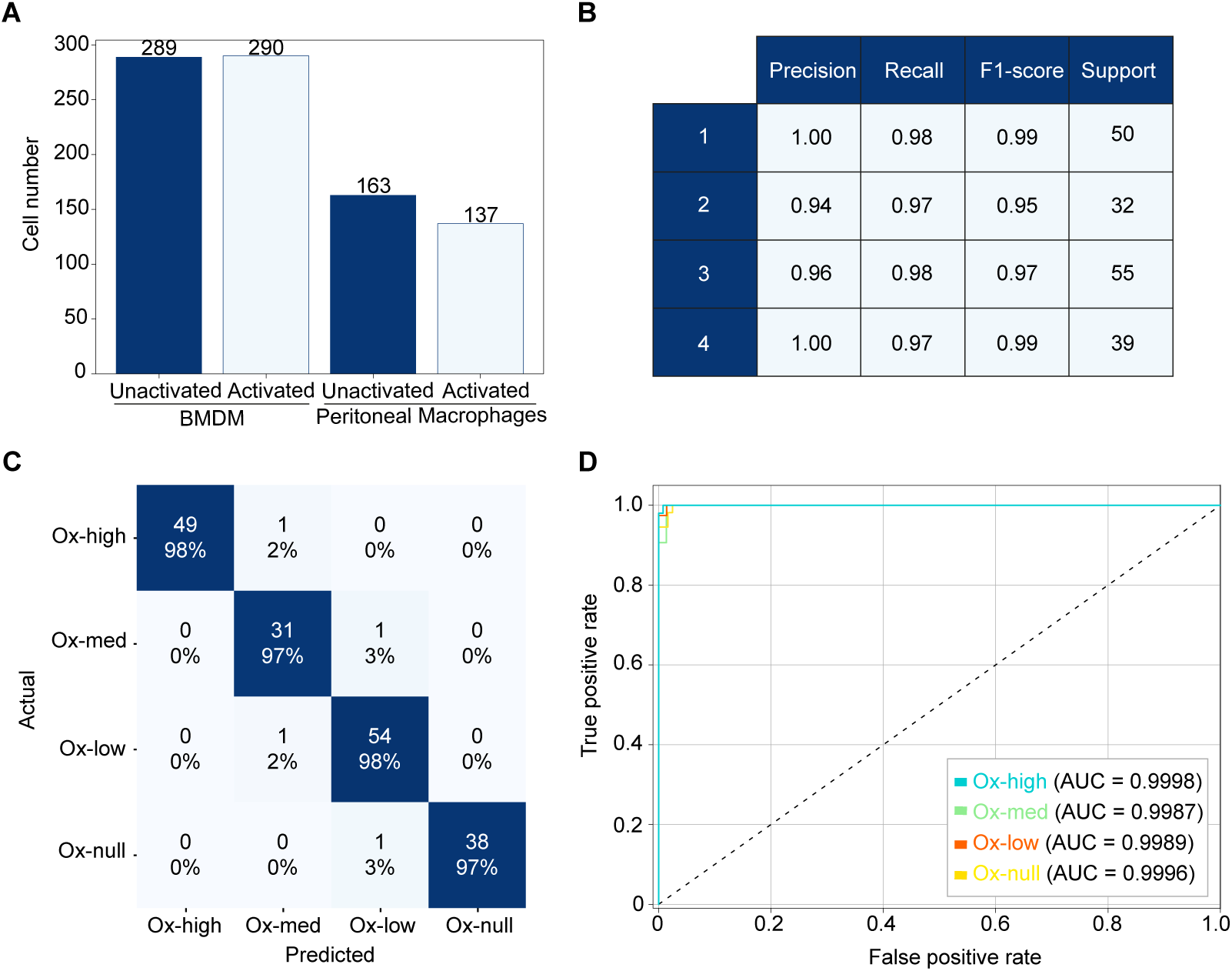
Performance of Met-Vision metabolic classifier. **A**) Distribution of the dataset including imaging of bone marrow derived macrophages and peritoneal macrophages treated or not with an inflammatory stimulus. **B**) Classification performance on the validation dataset. **C**) Confusion matrix of the predicted vs true labels for each class. **D**) AUCs of the Ox-high, Ox-med, Ox-low and Ox-null classes in the validation set.

**Figure S2:**
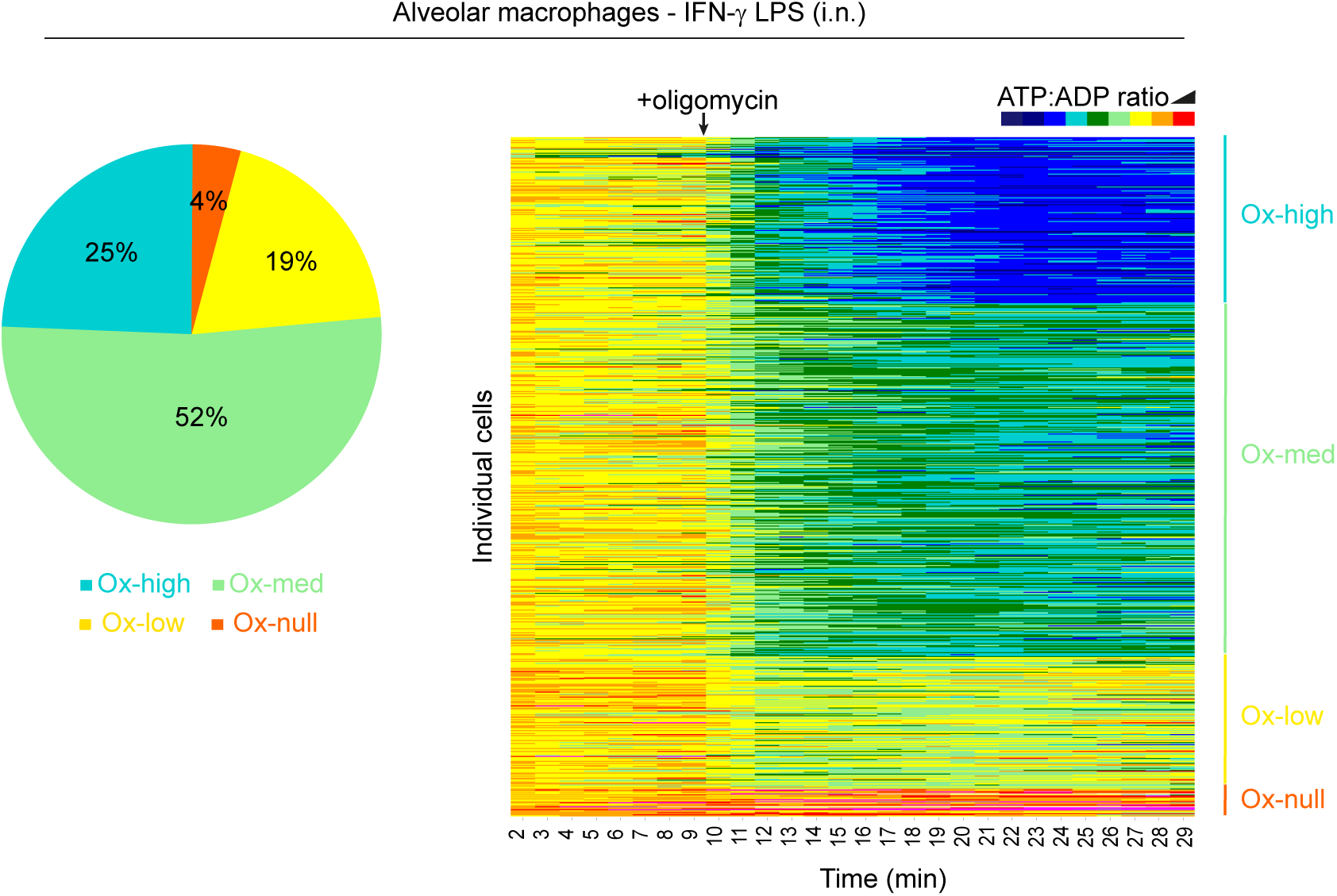
Distribution of macrophage energy metabolic profiles in alveolar macrophages upon instillation of LPS and IFN-γ. Alveolar macrophages were isolated from SPICY mice that were subjected to LPS+IFN-γ instillation two days before. Heatmaps and repartition of metabolic profiles within alveolar macrophages upon inflammation mediated by LPS+IFN-γ instillation.

## Movie legends

**Movie 1: Visualization of energy metabolism in single BMDMs**

The Met-Vision pipeline was applied to BMDMs from SPICY mice at steady-state or after activation with LPS and IFN-γ. After 10 min, oligomycin is added and imaging is continued. 64 macrophages are shown for each condition.

**Movie 2: Visualization of energy metabolism in single peritoneal macrophages**

The Met-Vision pipeline was applied to *ex vivo* isolated peritoneal macrophages from SPICY mice injected with PBS or LPS+IFN-γ two days before. After 10 min, oligomycin is added and imaging is continued. 64 macrophages are shown for each condition.

**Movie 3: Visualization of energy metabolism in single alveolar macrophages**

The Met-Vision pipeline was applied to *ex vivo* isolated alveolar macrophages from SPICY mice injected with PBS or LPS+IFN-γ two days before. After 10 min, oligomycin is added and imaging is continued. 64 macrophages are shown for each condition.

